# The Alzheimer’s disease risk gene CD2AP functions in dendritic spines by remodelling F-actin

**DOI:** 10.1101/2023.08.31.555707

**Authors:** Farzaneh S. Mirfakhar, Jorge Castanheira, Raquel Domingues, José S. Ramalho, Cláudia Guimas Almeida

**Affiliations:** iNOVA4Health, NOVA Medical School, Universidade Nova de Lisboa, 1169-056 Lisboa, Portugal; Department of Psychiatry, Washington University School of Medicine, St. Louis, MO 63110, USA

**Author notes:** These authors contributed equally (Farzaneh S. Mirfakhar, Jorge Castanheira).

**Keywords:** Actin-binding protein, late-onset Alzheimer’s disease, spines, synapse

## Abstract

CD2AP was identified as a genetic risk factor for late-onset Alzheimer’s disease (LOAD). However, how CD2AP contributes to LOAD synaptic dysfunction underlying AD memory deficits is unclear. We have shown that CD2AP loss-of-function increases β-amyloid (Aβ) endocytic production, but whether it contributes to synapse dysfunction is unknown. Because CD2AP is an actin-binding protein, it may also function in F-actin-rich dendritic spines, the excitatory postsynaptic compartment. Here, we demonstrate that CD2AP colocalises with F-actin in dendritic spines. Cell-autonomous depletion of CD2AP specifically reduces spine density and volume, with a functional decrease in synapse formation and neuronal network activity. Post-synaptic reexpression of CD2AP but not blocking Aβ-production is sufficient to rescue spine density. CD2AP overexpression increases spine density, volume, and synapse formation, while a rare LOAD CD2AP mutation induces aberrant F-actin spine-like protrusions without synapses. CD2AP controls postsynaptic actin turnover, with the LOAD mutation in CD2AP decreasing F-actin dynamicity. Our data support that CD2AP risk variants could contribute to LOAD synapse dysfunction by disrupting spine formation and growth by deregulating actin dynamics.

**Graphical abstract:** 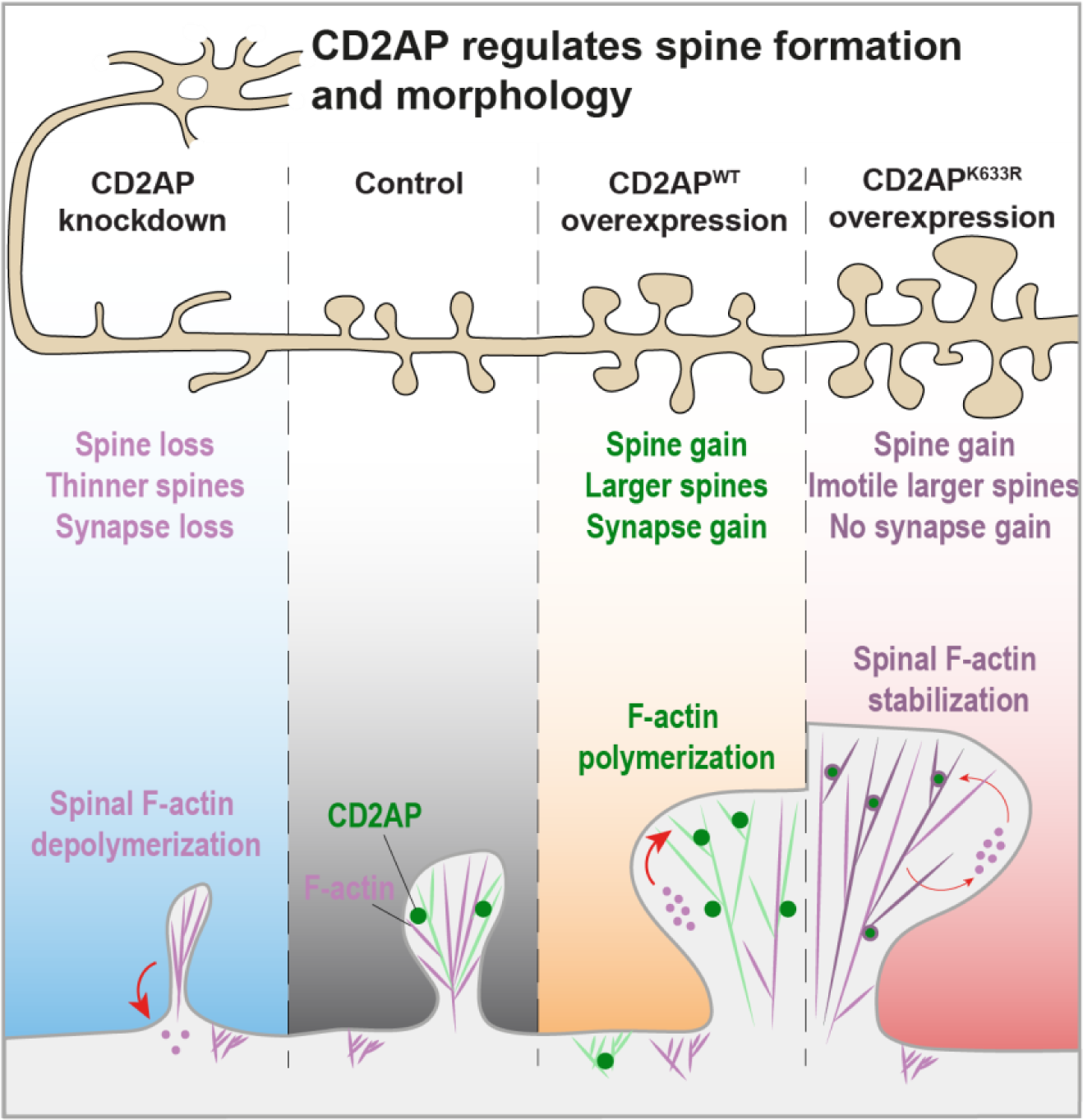

## Introduction

Late-onset Alzheimer’s disease (LOAD) is a devastating neurodegenerative disease that affects 1 in 9 people aged 65, increasing with aging (Alzheimer’s Association, 2023). Unfortunately, no effective treatment is available. A better understanding of the causal mechanisms may offer novel therapeutic strategies. The loss of neurones accompanying the pathology that defines AD, β-amyloid (Aβ) plaques, and tau neurofibrillary tangles is likely irreversible and untreatable. In contrast, early synapse dysfunction, which precedes neurodegeneration, may be modifiable. We believe it is necessary to dissect the causal mechanisms of synaptic dysfunction of LOAD to identify novel promising therapeutic targets.

*CD2AP* gene encoding for CD2-associated protein (CD2AP) was identified as a putative risk factor by genome-wide screens of thousands of LOAD patients (Hollingworth *et al.,* 2011; Naj *et al.,* 2011), confirmed in meta-analysis studies (Chen *et al.,* 2015; Gao *et al.,* 2022; Kamboh *et al.,* 2012; Kunkle *et al.,* 2019). While GWAS identified frequent noncoding variants, whole gene sequencing identified a pathological variant that introduces a mutation in CD2AP (Lys633Arg or K633R) highly associated with AD (Vardarajan *et al.,* 2015), but with unclear biological significance. The level of CD2AP brain expression in patients carrying CD2AP variants is unknown but was reduced in the blood of LOAD patients (Tao *et al.,* 2017) and accumulated in advanced AD (Camacho *et al.,* 2022). The CD2AP susceptibility loci correlate with the burden of neuritic plaques in AD patients (Shulman *et al.,* 2013).

CD2AP is an 80 kDa ubiquitously expressed scaffolding protein that belongs to the CIN85/CD2AP protein family, with three SH3 N-terminal domains, a proline-rich domain, and a coiled-coil c-terminal domain with actin-binding sites (Bruck *et al.,* 2006; Cormont *et al.,* 2003; Rouka *et al.,* 2015). Initially identified in T cells, CD2AP is important in kidney glomeruli cells, and its loss of function is related to proteinuria and renal dysfunction (Kim *et al.,* 2003; Shih *et al.,* 1999). CD2AP anchors slit diaphragm proteins to the actin cytoskeleton in glomeruli (Li *et al.,* 2000; Shih *et al.,* 2001); controls the polymerisation and stability of F-actin in undifferentiated cells (Tang & Brieher, 2013; Welsch *et al.,* 2001); interacts with F-actin, cortactin, or capping protein (CP) (Lynch *et al.,* 2003; Zhao *et al.,* 2013); and associates with membranes for endocytosis and endosomal maturation (Furusawa *et al.,* 2019; Gauthier *et al.,* 2007; Monzo *et al.,* 2005; Tolvanen *et al.,* 2015; Ubelmann *et al.,* 2017).

CD2AP is expressed in neurones (Li *et al.,* 2000; Ubelmann *et al.,* 2017), more in dendrites than in axons in mature neurones (Ubelmann *et al.,* 2017), consistent with postsynaptic function. CD2AP may also contribute to axonal neurite extension (Harrison *et al.,* 2016).

CD2AP was first associated with an AD mechanism by modulating tau toxicity in a Drosophila model of AD (Shulman *et al.,* 2014) and then to Aβ production but not to Aβ deposition in an amyloidosis model (PS1APP mice) (Liao *et al.,* 2015). We established that CD2AP loss of function, induced by shRNA-mediated depletion, disrupts the endocytic trafficking of Aβ precursor protein (APP), increasing its processing and Aβ generation at maturing early endosomes leading to intraneuronal Aβ accumulation in dendrites (Ubelmann *et al.,* 2017). However, it is unclear whether this increase in intraneuronal Aβ can cause synapse dysfunction in LOAD, as we showed in early-onset familial AD (eFAD) (Almeida *et al.,* 2005; Snyder *et al.,* 2005; Takahashi *et al.,* 2004).

CD2AP, like other LOAD risk genes, may cause synaptic dysfunction upstream Aβ production (Perdigão *et al.,* 2020). Since CD2AP is an actin regulator and the actin cytoskeleton is essential for spines that hold excitatory synapses (Hotulainen & Hoogenraad, 2010), we hypothesised that CD2AP might function at spines and thus impact synapses. Interestingly, the loss of Cindr, the Drosophila homolog of CD2AP, impaired neuromuscular synapses by interfering with synapse maturation (Ojelade *et al.,* 2019) supports our hypothesis.

Here, we investigate CD2AP synaptic function using knockdown and overexpression approaches in primary cultures of mouse cortical neurones and determine that CD2AP plays a direct role in spine density and morphology, modulating synaptic contacts. Notably, we found that a LOAD-associated mutation in CD2AP confers an aberrant gain of function in spines. Mechanistically, we established that CD2AP functions in the spines through spinal F-actin, with a minor contribution from Aβ production. Therefore, CD2AP risk variants can contribute to LOAD synapse dysfunction by deregulating spinal F-actin independently of Aβ.

## Results

### A pool of CD2AP localises in the spines

Previously, we localised CD2AP in dendritic endosomes relevant to APP endocytic trafficking in cortical mouse neurones (Ubelmann *et al.,* 2017). To determine if CD2AP localised to synapses, we started by analyzing CD2AP distribution in synaptic fractions of the mouse adult brain (Fig. 1A). We found CD2AP present in the postsynaptic fraction (P3) positive for postsynaptic density protein-95 (PSD-95) and negative for synapsin but not in the presynaptic fraction (S3) (Fig. 1A). Next, we investigated whether CD2AP was present in the spines. We detected CD2AP puncta in most spines. Interestingly, we frequently observed a punctum of CD2AP in the tip of the dendritic spines. We analysed the distribution of CD2AP in dendrites of mature cortical mouse neurones expressing mCherry to allow the morphological identification of dendritic spines. (Fig. 1B). We quantified the fraction of dendritic CD2AP or mCherry present in the spines and found that 34 % of dendritic CD2AP, more than soluble mCherry (27 %), localises to the spines of mature neurones (15 days *in vitro* (DIV) in BrainPhys media, or 21 DIV in Neurobasal media) (Fig. 1C). Furthermore, we measured CD2AP levels during the differentiation and synaptic maturation of primary mouse neurones cultured in BrainPhys media for 5, 11 and 15 DIV (Fig. 1D). CD2AP increased 3-fold from 5 to 15 DIV, similarly to PSD-95, the postsynaptic marker (Fig. 1E), supporting that CD2AP is expressed primarily in synaptically mature neurones and suggests a synaptic function for CD2AP. We also analysed the colocalisation of CD2AP with PSD-95 in neurones expressing GFP to identify dendritic spines (Fig. 1F) morphologically. Quantification revealed that 31 % of CD2AP puncta colocalised with PSD-95, which was equal to the % of PSD-95 puncta that colocalised with CD2AP (Fig. 1G), indicating that CD2AP is present in one-third of spines. In the 3D reconstruction (IMARIS) of a GFP-filled spine, the location of a CD2AP punctum at the tip of a spine, positioned laterally to the PSD-95 punctum, is highlighted (Fig. 1H). Endocytic zones are also lateral to the PSD (Lu *et al.,* 2007). Since we showed that CD2AP also localises to early endosomes, we include an example of a spine where a CD2AP punctum is laterally associated with the early endosome marker EEA1 to the F-actin-rich spine head (Fig. 1I). Furthermore, we imaged the movement of CD2AP-GFP puncta in a spine in mature live neurones (15 DIV) (Fig. 1G, supplementary movie 1).

**Figure 1.**
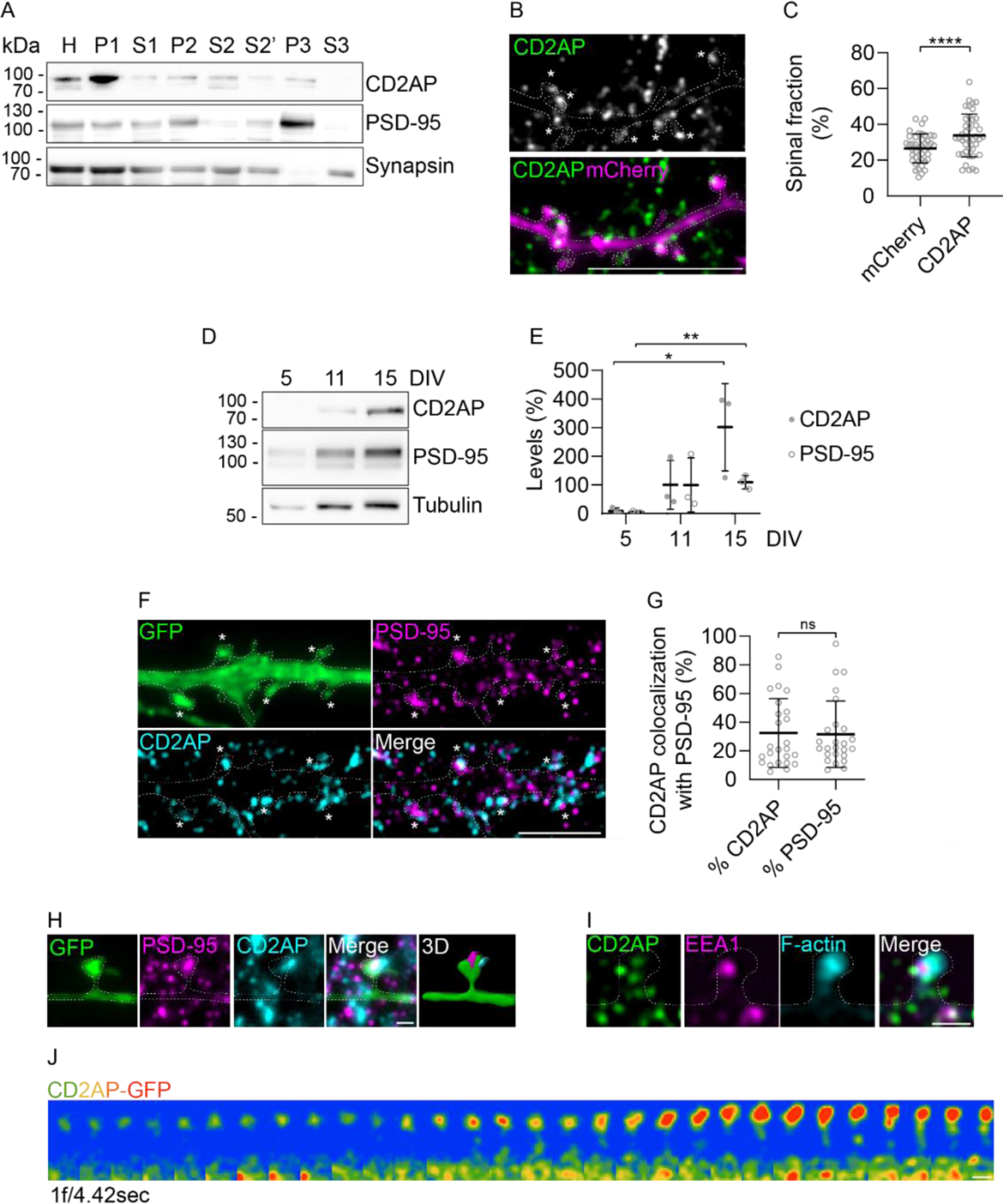
CD2AP is present in spines. **(A)** Western blot analysis of CD2AP, PSD-95, and synapsin in postnuclear supernatant (S1), crude synaptosomal fraction (P2), crude synaptic vesicle fraction (S3), and synaptosomal membrane fraction (P3) of the 6-month-old brain. Notably, CD2AP is present in the PSD-95 positive fraction P3 and not in the synaptic positive S3 fraction, supporting a postsynaptic location of CD2AP. **(B)** CD2AP (green) endogenous localisation in Cherry (magenta)-expressing dendrites Scale bar: 10 µm. **(C)** Quantification of the CD2AP and Cherry spinal enrichment, the fraction of dendritic signal in spines (n=4, N_Cherry_=42, N_CD2AP_=42 dendrites). **(D)** Western blot of CD2AP, PSD-95 and tubulin in primary neurones after 5, 11, and 15 DIV. **(E)** Quantification of CD2AP and PSD95 levels normalised to 11 DIV primary neurones (n=3). **(F)** Colocalisation of CD2AP (cyan) and PSD95 (magenta) in GFP-expressing dendritic spines. Scale bar: 5 µm. **(G)** Quantification of CD2AP colocalisation with PSD95 as a percentage of CD2AP and PSD-95 (n=2, N_CD2AP_=25, N_PSD-95_=25 dendrites,). **(H)** CD2AP (cyan) localisation lateral to PSD-95 (magenta) in the GFP-expressing spine (green). Scale bar: 1 µm. **(I)** CD2AP (green) colocalisation with EEA1 (magenta) and F-actin (cyan) at the spine. Scale bar: 1 µm. **(J)** CD2AP-GFP (red) enrichment at the spine head was recorded by time-lapse airy-scan confocal for 13 s (1 frame per 4.42 s). Scale bar: 0.5 µm.

Together, these data demonstrate that a pool of CD2AP is localised to the spines in mature primary cortical neurones.

### The removal of CD2AP reduces spine density and synapses

The localisation of CD2AP in spines led us to investigate whether CD2AP has synaptic function using a knockdown approach. To knock down CD2AP, we infected neurones with lentivirus expressing GFP and different sequences of small hairpin interfering RNA against CD2AP (shCD2AP-1 and shCD2AP-2)(Zhao *et al.,* 2013) or a nontargeting shRNA sequence (shControl). We confirmed the CD2AP knockdown by western blot using a mouse neuroblastoma cell line, Neuro2a cells and mouse fibroblasts (Fig. S1A, B).

To determine the impact of CD2AP knockdown on spines, we analysed spines, morphologically identified using the expression of GFP in shCD2AP- and shControl-treated neurones by epifluorescence microscopy and 3D reconstruction using IMARIS. We observed fewer and thinner spines after CD2AP knockdown (Fig. 2A). Quantifying spine density, the number of spines per dendritic length (10 μm), revealed that it was reduced by 55 % and 60 % with shCD2AP-1 and shCD2AP-2 treatment, respectively (Fig. 2B). Furthermore, the volume of the remaining spines decreased by almost 30 % after treatment with shCD2AP-1 and shCD2AP-2 (Fig. 2C).

**Figure 2.**
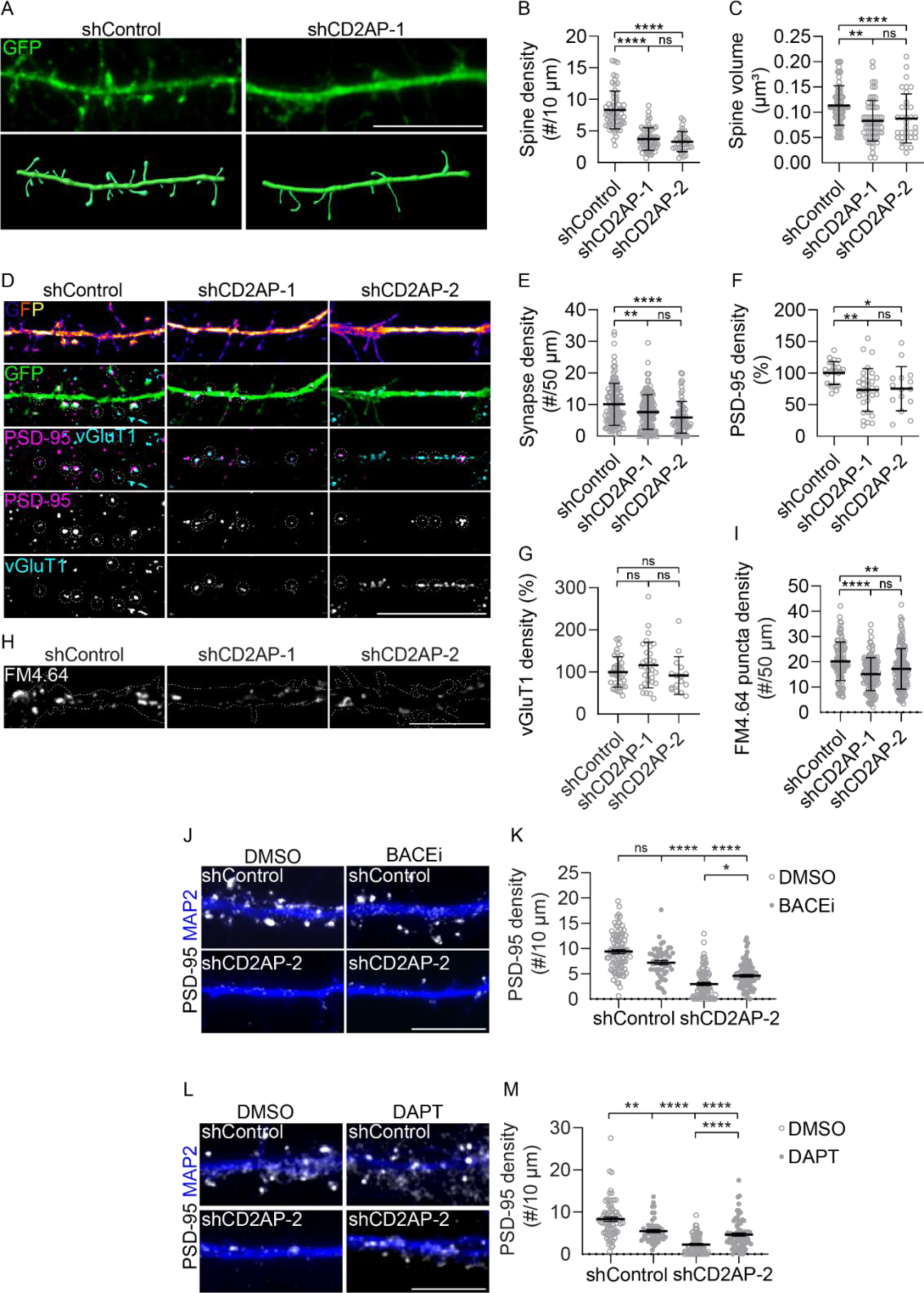
CD2AP knockdown reduces spine density and size and affects synapses. Primary neurones were treated with shControl, shCD2AP-1, or shCD2AP-2 as indicated. **(A)** Representative images of dendrites of neurones treated with shControl and shCD2AP-1 expressing GFP (green). Below is a 3D reconstruction using IMARIS. Scale bar: 10 µm. **(B)** Quantification of dendritic spine density (n = 5, N_shControl_ = 63, N_shCD2AP-1_ = 61, N_shCD2AP-2_ = 34 dendrites). **(C)** Quantification of dendritic spine volume based on GFP volume marker (n=5, N^shControl^=64, N_shCD2AP-1_=61, N_shCD2AP-2_=34 dendrites). **(D)** Synapses identified by PSD-95 (magenta, grey) and vGluT1 (cyan, grey) colocalisation in neurones expressing GFP (Fire LUT, green). Scale bar: 10 µm. **(E)** Quantification of the synapse density (n = 5, N_shControl_ l = 116, N_shCD2AP-1_ = 130, NshCD2AP-2 = 72 neurones). **(F)** Quantification of PSD-95 density (n = 5, N_shControl_ = 25, N_shCD2AP-1_= 31, N_shCD2AP-2_ = 15 neurones). **(G)** Quantification of vGluT1 density (n=5, N_shControl_=36, N_shCD2AP-1_=30, N_shCD2AP-2_=16 neurones). **(H)** Active synapses labelled with FM4.64 (grey) of neurones upon high-potassium-mediated depolarisation. Scale bar: 10 µm. **(I)** Quantification of FM4.64 puncta density (n=3, N_shControl_=110, N_shCD2AP-1_=123, N_shCD2AP-2_=151 neurites). **(J)** Spines identified with PSD-95 (grey) in dendrites labelled with MAP2 (blue) after treatment with BACE inhibitor (BACEi) or DMSO. Scale bar: 10 µm. **(K)** Quantification of PSD-95 density in dendrites of neurones after treatment with BACE inhibitor (BACEi) or DMSO (n=3, N_shControl+DMSO_=92, N_shCD2AP+DMSO_ =90, N_shControl+BACEi_=51, N_shCD2AP+BACEi_ =85 dendrites). **(L)** Spines identified with PSD-95 (grey) in dendrites labelled with MAP2 (blue) after treatment with γ-secretase inhibitor (DAPT) or DMSO. Scale bar: 10 µm. **(M)** Quantification of PSD-95 density after treatment with γ-secretase inhibitor (DAPT) or DMSO (n=3, N_shControl+DMSO_=74, N_shCD2AP+DMSO_ =81, N_shControl+DAPT_=56, N_shCD2AP+DAPT_ =79 dendrites). Data are presented as mean ± SD. \**P*<0.05, \*\**P*<0.01, \*\*\**P*<0.001, \*\*\*\**P*<0.0001; ns, not significant.

To assess whether CD2AP-dependent reduction in spines affected synapses, we performed immunostaining for vGluT1 and PSD-95 to identify pre- and postsynaptic compartments, respectively, of excitatory synapses. In dendrites depleted for CD2AP, there was a reduction in PSD-95 and vGluT1 puncta (Fig. 2D). We automatically quantified their colocalisation as a proxy for synapses. The synapse density along dendrites was reduced by 24 % and 41 % with treatment with shCD2AP-1 and shCD2AP-2, respectively (Fig. 2E). This synapse density reduction seems to be mainly accounted by the reduction in PSD-95 puncta density with shCD2AP-1 and shCD2AP-2 treatment (25 %) (Fig. 2F) since the vGluT1 puncta density remained unchanged (Fig. 2G). We measured the density of active synapses after shCD2AP treatment by monitoring the density of FM4.64 labelled presynaptic puncta after induction of high potassium-induced neuronal depolarisation (90 s) (Fig. 2H). The FM4.64 puncta density was significantly reduced by 25 % and 15 % with shCD2AP-1 and 2, respectively (Fig. 2I). These results support synapse reduction due to the lack of functional spines in the absence of CD2AP.

Since we and others showed that CD2AP knockdown increases intraneuronal Aβ42 (Ubelmann *et al.,* 2017; Liao *et al.,* 2015) and that Aβ in early-onset transgenic mouse primary cortical neurones reduces spines (Almeida *et al.,* 2005), we investigated whether reduction in spine density would be rescued by blocking Aβ production as previously (Burrinha *et al.,* 2021). We measured the density of PSD-95 puncta in shCD2AP neurones treated with the BACE inhibitor (BACEi) and the gamma-secretase inhibitor (DAPT) (Fig. 2J-M). Treatment with BACEi and DAPT increased PSD-95 puncta density in neurones treated with shCD2AP-2, but remained significantly reduced compared to shControl (Fig. 2J-M). Notably, DAPT, but not BACEi, treatment reduced PSD-95 density in shControl neurones, supporting that γ-secretase has synaptic relevant substrates (Bittner *et al.,* 2009; Barthet *et al.,* 2018; Servián-Morilla *et al.,* 2018). These results support the notion that Aβ production contributes but does not fully account for spine loss induced by CD2AP knockdown.

Thus, we discovered that CD2AP depletion reduces synapses by affecting the spines in part independently of Aβ production.

### CD2AP knockdown impacts neuronal network activity

To investigate the impact of CD2AP knockdown on basal neuronal activity, we performed multielectrode array (MEA) recordings on primary neurones cultured in BrainPhys for 15 DIV. A representative whole MEA (4096 electrodes) shows bright red/yellow/white active electrodes in the shControl-treated neurones, consistent with spontaneous neuronal activity (Fig. 3A). The neurones showed less bright red/yellow/white electrodes. We quantified the percentage of active electrodes displaying neuronal activity that registered more than three spikes per s and found, on average, less active electrodes in shCD2AP-2 neurones (2.9 %) and a tendency to decrease in shCD2AP-1 neurones compared to shControl neurones (6 %) (Fig. 3B), supporting that CD2AP depletion suppresses neuronal activity.

**Figure 3.**
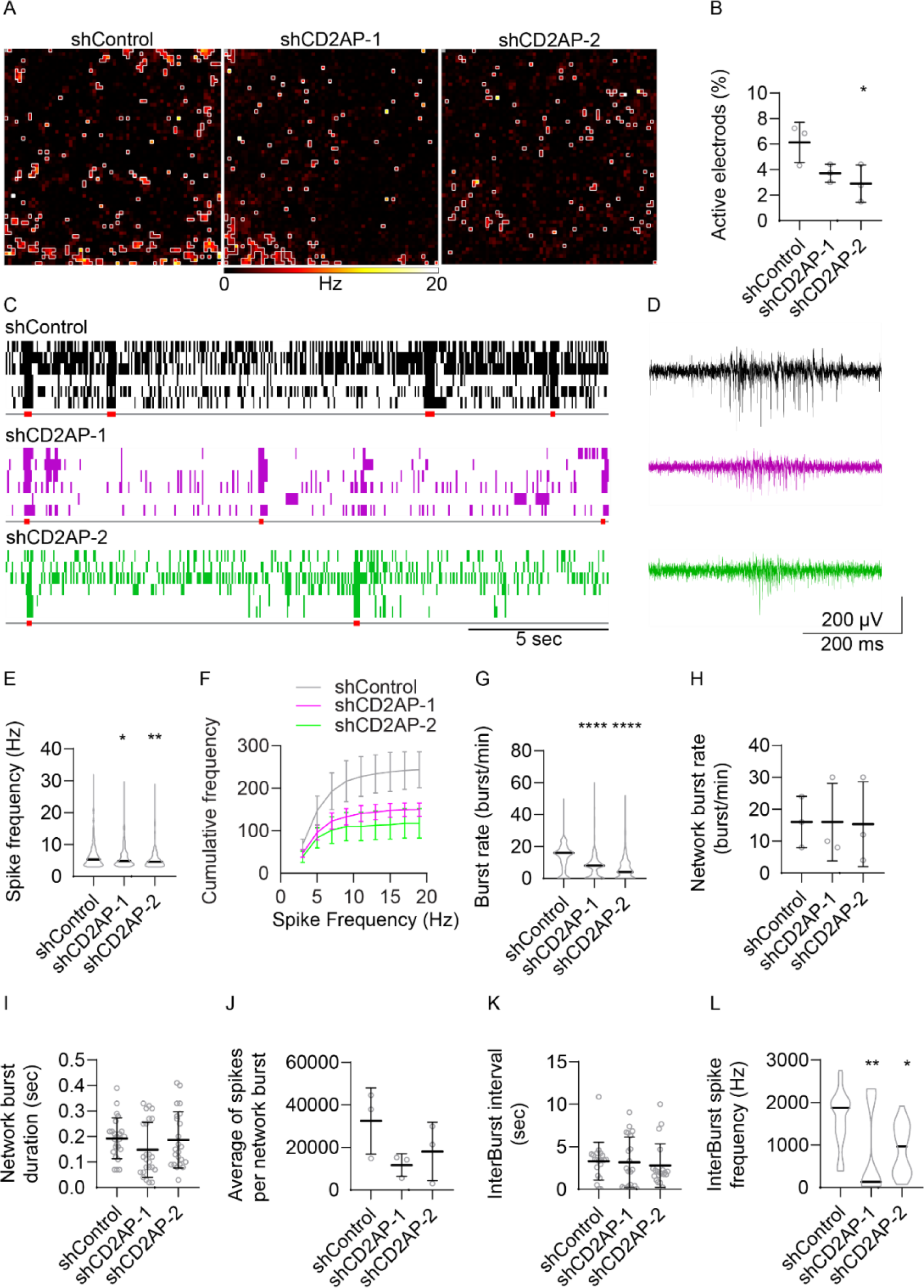
CD2AP knockdown reduces neuronal activity. Primary neurones treated with shControl, shCD2AP-1, or shCD2AP-2 and basal activity were recorded with an MEA of 4096 electrodes. **(A)** Representative heatmaps of the spiking activity of active electrodes with more than three spikes per s are outlined in white. **(B)** Percentage of active electrodes (n=3). **(C)** Representative raster lines showing spiking and burst activity of shControl (black), shCD2AP-1 (magenta) and shCD2AP-2 (green) treated neurones. Network bursts are identified below with red squares. **(D)** Representative traces showing burst spiking activity of shControl (black), shCD2AP-1 (magenta) and shCD2AP-2 (green) neurones. **(E)** Average spike frequency (Hz) of shControl (grey), shCD2AP-1 (magenta) and shCD2AP-2 (green) neurones (n=3, N_shControl_=751, N_shCD2AP-1_=452, N_shCD2AP-2_=356 electrodes). **(F)** Cumulative spike frequency of shControl (grey), shCD2AP-1 (magenta) and shCD2AP-2 (green) (n=3, N_shControl_=751, N_shCD2AP-1_=453, N_shCD2AP-2_=356). **(G)** Burst rate per minute (n=3, N_shControl_=753, N_shCD2AP-1_=456. N_shCD2AP-2_=355 bursts). **(H)** Network burst rate per minute (n=3). **(I)** Network burst duration in s (n=3, N_shControl_=24, N_shCD2AP-1_=29, N_shCD2AP-2_=22 bursts). **(J)** Average spikes per network burst (n=3). **(K)** Interburst interval (n=3, N_shControl_=21, N_shCD2AP-1_=26, N_shCD2AP-2_=19 intervals). **(L)** Interburst spike frequency (n=3, N_shControl_=24, N_shCD2AP-1_=29, N_shCD2AP-2_=22). Data are presented as mean ± SD. \**P*<0.05, \*\**P*<0.01, \*\*\*\**P*<0.0001.

The raster profiles show the lower spiking and burst activity of shCD2AP-1 and shCD2AP-2 neurones compared to shControl neurones (Fig. 3C). The spike frequency on each active electrode, as in the representative traces (Fig. 3D), was reduced in shCD2AP-1 neurones (6.0 Hz) and shCD2AP-2 neurones (5.9 Hz) compared to shControl neurones (6.7 Hz) (Fig. 3E). The cumulative frequency of spiking neurones showed a lower frequency of high spiking neurones after shCD2AP-1 and -2 treatment than in control neurones, suggesting that depleting CD2AP mitigates neural excitability (Fig. 3F).

In the raster profiles, we also observed synchronised bursts of activity in the neuronal network. By calculating the burst rate (burst/min) in each active electrode, we found a lower burst frequency in shCD2AP-1 (8 burst/min) and shCD2AP-2 neurones (7 burst/min) compared to shControl neurones (13 burst/min). Although no significant changes were observed in network burst rate (Fig. 3H), network burst duration (Fig. 3I), average of spikes per network burst (Fig. 3J) and interburst interval (Fig. 3J) with the knockdown of CD2AP, the interburst spike frequency reduced from 717.8 Hz in shControl neurones to 269.7 Hz and 462.8 Hz, in shCD2AP-1 and shCD2AP-2 treated neurones, respectively (Fig. 3L). These results indicate that the basic structure of the burst events and the time between them remained stable despite the overall reduction in frequency, consistent with a decrease in the synchronised activity of the neuronal network in the absence of CD2AP.

The findings suggest that the impact on neural activity is complex and may involve changes in network synchrony and excitability. Further investigation is warranted to understand the underlying mechanisms.

### CD2AP mutant increases spine density more than CD2AP wild-type overexpression

After establishing that CD2AP knockdown reduced spines and synaptic activity, we investigated the impact on synapses of the overexpression of CD2AP and of the K633R coding mutation in CD2AP identified in LOAD patients and associated with a higher risk of LOAD (Vardarajan *et al.,* 2015). We transfected primary neurones with CD2AP^WT^-GFP CD2AP^K663R^-GFP and the control GFP plasmid at 7 DIV and evaluated spinal and synaptic alterations at 15 DIV (Fig. S2 A). We mutagenised CD2AP^WT^-GFP to obtain CD2AP^K633R^-GFP (Fig S2 B).

CD2AP^WT^ and CD2AP^K633R^ expression were evident in dendrites and dendritic protrusions like spines (Fig. 4A). We noted that overexpression of CD2AP^WT^ led to an increase in the number of primary dendrites, their length, and the number of intersections, unaltered by the LOAD mutation (Fig. S2 D-H).

**Figure 4.**
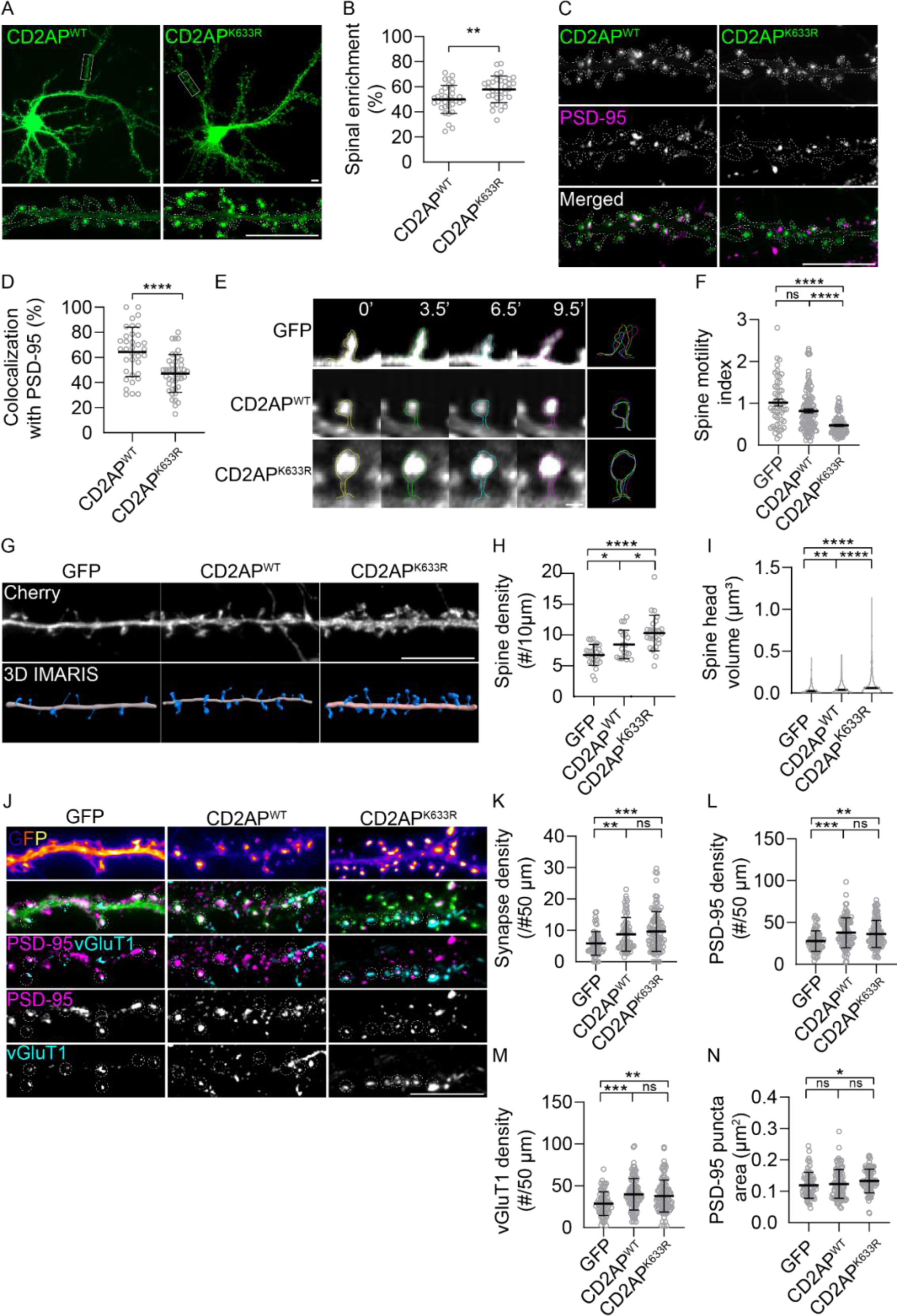
The CD2AP LOAD mutation increases spine density and volume but not synapse density induced by CD2AP overexpression. Primary neurones overexpressing CD2AP^WT^-GFP, CD2AP^K633R^-GFP, or GFP as control as indicated. **(A)** Representative images of neurones overexpressing CD2AP^WT^-GFP and CD2AP^K633R^-GFP (green). White squares indicate dendritic segments magnified below. Scale bar: 10 µm. **(B)** Quantification of the CD2AP^WT^ and CD2AP^K633R^ spinal enrichment, the fraction of dendritic signal in spines (n=5, N ^WT^=33, N ^K633R^=32). **(C)** CD2AP^WT^ or CD2AP^K633R^ (grey, green) colocalisation with PSD-95 (grey, magenta) alone and merged. Scale bar: 10 µm. **(D)** Quantification of the colocalisation of CD2AP^WT^ or CD2AP^K633R^ with PSD95 (% of PSD-95) (n=3, N ^WT^=35, N ^K633R^=42 dendrites). **(E)** Representative time series of spines expressing GFP, CD2AP^WT,^ or CD2AP^K633R^ at 0, 3.5, 6.5, and 9.5 min, outlined in yellow, green, blue, and magenta, respectively. The panel on the right shows superimposed spine outlines to reveal spine motility over time. Scale Bar: 1 µm. **(F)** Quantification of spine motility index (n=2, N_GFP_=49, N_CD2AP_^WT^=112, N_CD2AP_^K633R^=92 spines). **(G)** Representative images of dendrites of neurones expressing with the volume marker Cherry (grey). Below is a 3D reconstruction using IMARIS. Scale bar: 10 µm. **(H)** Quantification of dendritic spine density (n=3, N_GFP_=35, N_CD2AP_^WT^=30, N_CD2AP_^K633R^=32 dendrites). **(I)** Quantification of spine head volume based on Cherry volume marker (n=3, N_GFP_=49, N ^WT^=52, N ^K633R^=79 spines). **(J)** Synapses identified by PSD-95 (magenta, grey) and vGluT1 (cyan, grey) colocalisation in GFP, CD2AP^WT,^ or CD2AP^K633R^ expressing neurones (fire LUT, green). Scale bar: 10 µm. **(K)** Quantification of synapse density (n=3, N_GFP_=66, N_CD2AP_^WT^=66, N_CD2AP_^K633R^=99 dendrites). **(L)** Quantification of PSD-95 (n=3, N_GFP_=70, N_CD2AP_^WT^=89, N_CD2AP_^K633R^=97 dendrites). **(M)** Quantification of vGluT1 density in GFP, CD2AP^WT,^ or CD2AP^K633R^ neurites (n=3, N_GFP_=71, N ^WT^=94, N_CD2AP_^K633R^=98 density). **(N)** Quantification of PSD-95 average area (n=3, N_GFP_=67, N ^WT^=93, N_CD2AP_^K633R^=91 dendrites). Data are presented as mean ± SD. \**P*<0.05, \*\**P*<0.01, \*\*\**P*<0.001, \*\*\*\**P*<0.0001; ns, not significant.

Interestingly, 50 % of CD2AP^WT^ and 58 % of CD2AP^K633R^ are in dendritic protrusion-like spines (Fig. 4A-B), representing an increase of 47 % and 71 % relative to endogenous CD2AP (34 %; Fig.1C), respectively. These data indicate that the LOAD mutation in CD2AP increases its expression in spines. We assessed whether CD2AP-positive protrusions were spines by labelling with PSD-95 (Fig. 4C). Colocalisation analysis revealed the percentage of PSD-95 puncta positive for CD2AP^K633R^ (47 %) inferior to CD2AP^WT^ (64 %), indicating that there are more spines positive for CD2AP^WT^ than for CD2AP^K633R^ (Fig.4D).

Dendritic spines are highly motile *in vitro* and *in vivo* (Lendvai *et al*., 2000; Tashiro & Yuste, 2004). This motility is likely related to extensions and retractions as if searching for presynaptic partners. Spine motility also allows for maturation-related morphological changes (Bonhoeffer & Yuste, 2002). Live neurones were imaged every 30 s for 10 minutes to examine rapid spine motility (Fig. 3E) (Tashiro & Yuste, 2004). We observed spines showing motility with expansions and contractions during imaging. We measured the spine area during the imaging and calculated the spine motility index to quantify the degree of spine motility (Dunaevsky *et al.,* 1999). The motility index was quantified by calculating the difference between the largest and smallest spine area, normalised to the average spine area during the movie. While CD2AP^WT^ tended to decrease spine motility, only the CD2AP^K633R^-positive spines significantly reduced motility by more than 50 % (Fig. 4F).

CD2AP^WT^ and CD2AP^K633R^-positive spines were denser and bulkier than the control (GFP) (Fig. 4G). We transfected primary neurones with CD2AP^WT^ or CD2AP^K633R^ and the volume marker mCherry for spine density and spine head volume analysis. Spines were manually identified and counted based on mCherry expression. Spine density increased (25 %) in neurones expressing CD2AP^WT^ (8.3 spines/10 µm) and increased by 53 % in neurones expressing CD2AP^K633^ (10.1 spines/10 µm) relative to control neurones (GFP) (6.6 spines/10 µm) (Fig. 4H).

Spine head volume was measured upon 3D reconstruction with IMARIS software based on mCherry expression. Overexpression of CD2AP^WT^ increased the spine head volume by 26 % (0.058 µm^3^) and the CD2AP^K633R^ overexpression increased 2.5 times the volume of the head of the spine (0.115 µm^3^) compared to control spines (0.046 µm^3^) (Fig. 4I). These results together with the negative impact of CD2AP knockdown on spines (Fig. 2A-C) indicate that CD2AP is a positive regulator of spine formation and growth and that the LOAD K633R mutation could enhance CD2AP spinal function.

To assess the impact of CD2AP^WT^ and CD2AP^K633R^ expression on excitatory synapses, we analysed vGluT1 and PSD-95 colocalisation (Fig. 4J). Synapse density increased with CD2AP^WT^ expression (Fig. 4K). As did PSD-95 and vGluT1 density increased upon CD2AP^WT^ expression (Fig. 4L-M). These results indicate that the postsynaptic expression of CD2AP^WT^ likely increased PSD-95, consistent with the induction of the formation and growth of mature spines. This postsynaptic expansion may translate into a presynaptic adaptation in the non-expressing presynaptic neurone since vGluT1 accompanied the increase in PSD-95. Unexpectedly, given the increase in spine density and volume induced by CD2AP^K633R^ expression, synapse density only increased with CD2AP^WT^ expression (Fig. 4K-M). In particular, PSD-95 area increased in neurones (Fig. 4N). The LOAD CD2AP^K633R^ did not enhance synapse density; instead, it interfered with the function of CD2AP.

These results and the negative impact of CD2AP knockdown on spines and synapses support that CD2AP has a postsynaptic function that impacts synapses.

### CD2AP controls spinal F-actin

We established that CD2AP functions in spine formation and growth and that a LOAD risk variant may interfere with its function. Since CD2AP is an F-actin binding protein and a regulator of actin dynamics, we investigated whether CD2AP regulates spines through F-actin.

We colocalised CD2AP with F-actin, labelled with phalloidin, and cortactin, a spinal F-actin regulator (Schnoor *et al.,* 2018). We observed F-actin foci present in most spines, while CD2AP, like cortactin, was detected in a subset of spines (Fig. 5A). In higher magnification, a partial overlap between CD2AP and cortactin puncta with F-actin foci in spine heads was detected (Fig. 5A). We found a 30 % colocalisation of CD2AP with F-actin and cortactin (Fig. 5B).

**Figure 5.**
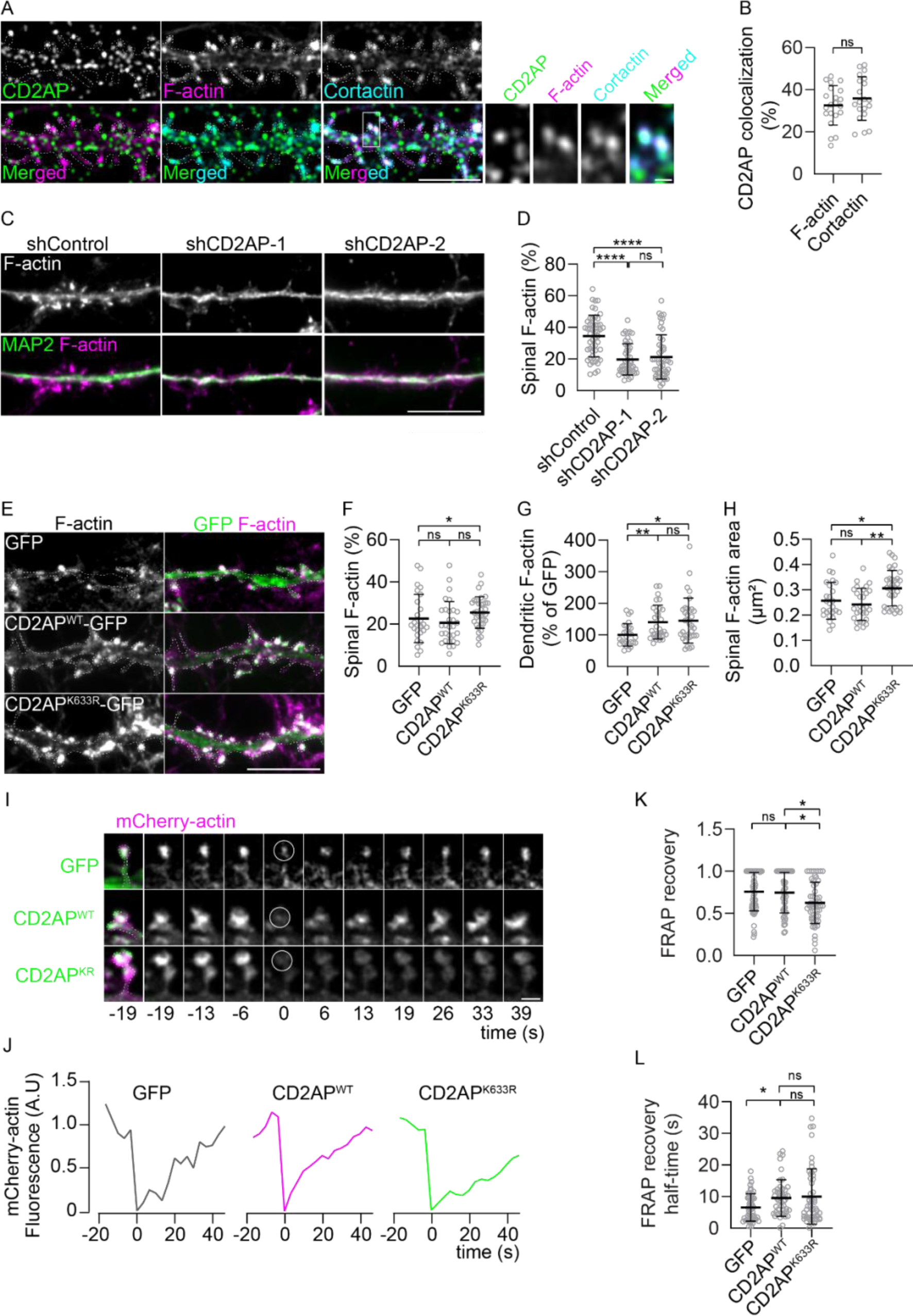
CD2AP modulates spinal F-actin. **(A)** CD2AP (green), F-actin (magenta), and cortactin (cyan) colocalisation in dendrites. Scale bar: 5 µm. The white rectangle indicates magnified spines on the right. Scale bar: 0.5 µm. **(B)** Quantification of CD2AP colocalisation with F-actin and cortactin (n=2, N=13-14 dendrites). **(C)** Spinal F-actin (grey; magenta) alone and colocalised with MAP2 (green) in dendrites of shControl, shCD2AP-1, or shCD2AP-2-treated neurones. Scale bar: 10 µm. **(D)** Quantification of spinal F-actin in dendrites of neurones treated with shControl, shCD2AP-1 or shCD2AP-2 (n=4, N_shControl_=50, N_shCD2AP-1_=48, N_shCD2AP-2_=49 dendrites). **(E)** Spinal F-actin (grey; magenta) in dendrites of neurones expressing GFP, CD2AP^WT^, and CD2AP^K633R^ (green). Scale bar: 10 µm. **(F)** Quantification of spinal F-actin in dendrites of neurones expressing GFP, CD2AP^WT,^ and CD2AP^K633R^ (n=3, N_GFP_=26, N_CD2AP_^WT^=30, N_CD2AP_^K633R^=34 dendrites). **(G)** Quantification of F-actin mean intensity in dendrites of neurones expressing GFP, CD2AP^WT^, and CD2AP^K633R^, normalised to GFP (n=3, N_GFP_=27, N_CD2AP_^WT^=30, N_CD2AP_^K633R^=34). **(H)** Quantification of spinal F-actin average area in dendrites of neurones expressing GFP, CD2AP^WT,^ and CD2AP^K633R^ (n=3, N_GFP_=26, N ^WT^=29, N_CD2AP_^K633R^=34 dendrites). **(I)** Representative mCherry-actin (magenta) in the spines of dendrites of neurones expressing GFP, CD2AP^WT,^ and CD2AP^K633R^ (green). The right panels correspond to frames at the indicated times of the corresponding FRAP experiment of mCherry-actin (grey). Scale bar: 1 µm. **(J)** Representative FRAP traces of normalised mCherry-actin fluorescence (A.U.) in the spines expressing GFP (grey), CD2AP^WT^ (magenta), and CD2AP^K633R^ (green) are shown in (I). **(K)** Quantification of mCherry-actin FRAP recovery (n=4, N_GFP_=55, N_CD2AP_^WT^=458, N_CD2AP_^K633R^=55 spines). **(L)** Quantification of mCherry-actin FRAP recovery half-time (n=4, N_GFP_=55, N ^WT^=458, N ^K633R^=55 spines). Data are presented as mean ± SD. \**P*<0.05, \*\*\*\**P*<0.0001; ns, not significant.

Next, we evaluated the impact of CD2AP knockdown on spinal F-actin. We observed less F-actin in the spines of shCD2AP-treated neurones (Fig. 5C). Quantification revealed that 34 % of dendritic F-actin is in the spines of neurones treated with shControl. In neurones treated with shCD2AP-1 and shCD2AP-2, the distribution of F-actin between the shaft and spines reduced to 19 % and 21 % of dendritic F-actin, respectively (Fig. 5D).

In CD2AP^WT^ expressing dendrites, the % of spinal F-actin was like in control dendrites (GFP) (Fig. 5E-F). In contrast, we found that F-actin mean intensity increased in CD2AP^WT^ expressing dendrites relative to control (Fig. 5E, G). Differently in spines expressing CD2AP^K633R^, F-actin increased (Fig. 5E-F), with puncta with a larger area (Fig. 4H). These results indicate that changes in F-actin likely mediate CD2AP-dependent spine expansion. Furthermore, the LOAD mutation in CD2AP may modify its function, inducing a build-up of F-actin in the spines.

To determine the impact of CD2AP^WT^ or CD2AP^K633R^ overexpression on actin turnover (Koskinen & Hotulainen, 2014), we measured the rate of actin filament assembly by co-expressing mCherry-actin (Koestler *et al.,* 2008) and performing fluorescence recovery after photobleaching (FRAP) imaging experiments (Fig. 5I, supplementary FRAP movies). Consistent with previous reports (Koskinen & Hotulainen, 2014), we observed that the fluorescence of mCherry-actin, after photobleaching correction and background subtraction, recovered in GFP expressing spines at 76% of the prebleached levels (Fig. 5I, K). In spines expressing, CD2AP^WT^ mCherry-actin recovered (75 %) similarly to control spines, suggesting that CD2AP^WT^ does not alter the stable fraction of F-actin (Fig. 5I, K). In contrast, mCherry-actin recovery in spines expressing CD2AP^K633R^ was significantly decreased (65 %) compared to spines-expressing GFP and CD2AP^WT^, suggesting that the LOAD mutation reduces the dynamic actin turnover by increasing the stable fraction of F-actin (Fig. 5I, K). Differently, the half-time of recovery increased in CD2AP^WT^-expressing spines (9.5 s), and tended to increase in CD2AP^K633R^-expressing spines (10.0 s) compared to GFP-expressing spines (6.5 s), indicative of an increase in the time needed for actin monomers bind to the polymerizing actin filaments, suggesting that CD2AP may sterically interfere with mCherry-actin binding to F-actin or cap the growing F-actin barbed end resulting in a less dynamic actin turnover in spines (Fig. 5 L). To correlate the rate of actin recovery with the head size of the spine, we categorised the spines as small (<0.2 um^2^), intermediate and large (>0.6 um^2^) (Fig. S3). The large GFP control spines tended to recover less, likely more stable, while the larger CD2AP^WT^ and CD2AP^K633R^ expressing spines tended to recover more than larger GFP expressing spines and smaller spines. Interestingly, the intermediate-size spines expressing CD2AP^K633R^ were significantly reduced compared to control or intermediate spines expressing CD2AP^WT^, supporting an increase in the stability of F-actin, especially in intermediate-size spines (Fig.S3).

### The CD2AP LOAD mutation does not rescue synapses or spinal F-actin

Finally, we investigated whether the loss of F-actin-positive spines induced by CD2AP depletion could be rescued by the reexpression of CD2AP^WT^ or CD2AP^K633R^ (Fig.6). We found that CD2AP^WT^ rescued the reduction in PSD-95 density induced by shCD2AP-1 and -2 to control levels, while in contrast the CD2AP^K633R^ was unable to increase PSD-95 density (Fig.6A, B). We also evaluated the impact on the PSD-95 area and observed a similar trend, with only the expression of CD2AP^WT^ significantly increasing the PSD-95 area compared to the neurones treated with shCD2AP-1 treated neurones (Fig.6C).

**Figure 6.**
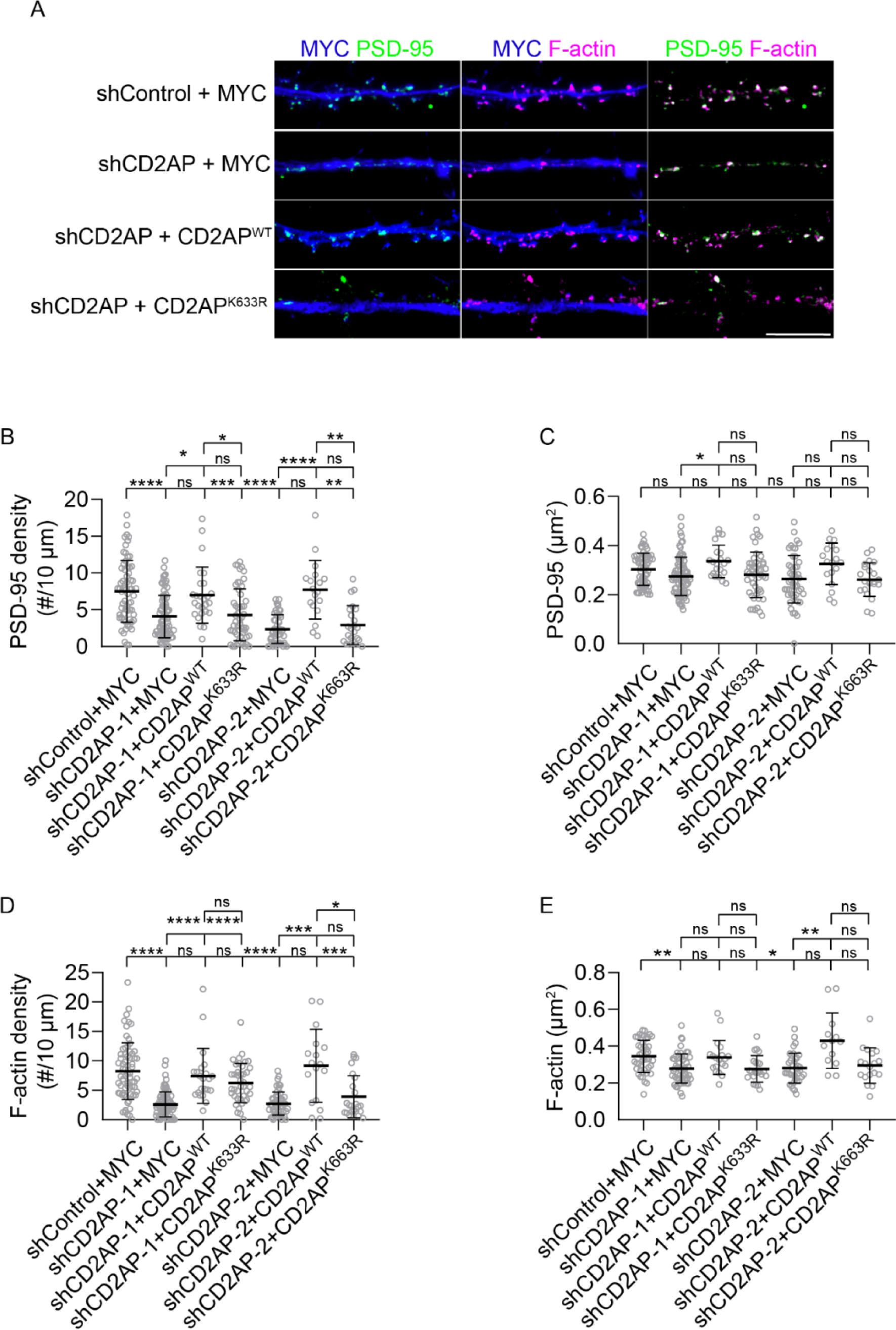
CD2AP wild-type but not mutant rescues spine density. shControl, shCD2AP-1, and shCD2AP-2 treated neurones expressing Myc, CD2AP^WT^-MYC, or CD2AP^K633R^-MYC. **(A)** PSD-95 (green), F-actin (magenta) in dendrites of shControl or shCD2AP-2-treated neurones expressing Myc, CD2AP^WT^, or CD2AP^K633R^ (blue). Scale bar: 10 µm. **(B)** Quantification of PSD-95 density (n=4, N_shControl+myc_=67, N_shCD2AP-1+myc_=83, N_shCD2AP-2+myc_=52, N_shCD2AP-2+WT_=18, N_shCD2AP-2+K633R_=23, N_shCD2AP-1+WT_=26, N_shCD2AP-2+K633R_=52 dendrites). **(C)** Quantification of PSD-95 puncta area (n=4, N_shControl+myc_=66, N_shCD2AP-1+myc_=82, N_shCD2AP-2+myc_=49, N_shCD2AP-2+WT_=18, N_shCD2AP-2+K633R_=23, N_shCD2AP-1+WT_=22, N_shCD2AP-1+K633R_=50 dendrites). **(D)** Quantification of F-actin puncta density (n=4, N_shControl+myc_=67, N_shCD2AP-1+myc_=83, N_shCD2AP-2+myc_=50, N_shCD2AP-2+WT_=17, N_shCD2AP-2+K633R_=24, N_shCD2AP-1+WT_=22, N_shCD2AP-1+K633R_=40 dendrites). **(E)** Quantification of F-actin puncta area (n=4, N_shControl+myc_=66, N_shCD2AP-1+myc_=82, N_shCD2AP-2+myc_=49, N_shCD2AP-2+WT_=18, N_shCD2AP-2+K633R_=23, N_shCD2AP-1+WT_=22, N_shCD2AP-1+K633R_=50 dendrites). Data are presented as mean ± SD. \**P*<0.05, \*\**P*<0.01, \*\*\**P*<0.001, \*\*\*\**P*<0.0001; ns, not significant.

We additionally analysed the density and area of F-actin puncta in dendrites. We found that CD2AP^WT^ but not CD2AP^K633R^ rescued F-actin density, similarly to PSD-95 (Fig. 6D). Concerning the F-actin area, the reduction was significant both in shCD2AP-1 and shCD2AP-2 treated neurones, and the rescue by CD2AP^WT^ reached significance in shCD2AP-2 treated neurones (Fig. 5E).

These results indicate that CD2AP controls synapses through spine formation and growth and that the LOAD mutation K633R causes loss of function, likely interfering with CD2AP control of spinal F-actin.

These results support a new specific function for CD2AP in controlling the balance between F-actin polymerisation and depolymerisation, the actin dynamics required for spine formation and growth. In addition, LOAD mutations may be associated with faulty F-actin dynamics that affects spine formation and growth, compromising synapses.

## Discussion

Spines depend on the protrusive force of actin polymerisation for filopodia formation and spine expansion by recruiting postsynaptic scaffolds, such as PSD-95, which allow the formation, plasticity, and strength of synapses. Actin-binding proteins regulate the dynamics of F-actin in spines (Bosch & Hayashi, 2012). Here, we discovered that the Alzheimer’s disease risk factor CD2AP, an actin-binding protein, is a postsynaptic protein. We localised CD2AP to spines and found that it is required for its formation and expansion, affecting the density of the synapses and the electrophysiological activity of the neurones. CD2AP likely functions in the spines via the actin cytoskeleton, with loss of spinal F-actin while overexpression increased it. Notably, the coding variant in CD2AP (K633R) associated with LOAD induces aberrant spine-like formation and expansion. Not being able to rescue the CD2AP knockdown synapse phenotype suggests that the LOAD mutation may cause loss of CD2AP function. The LOAD mutation decreased the FRAP actin recovery, potentially interfering with the depolymerization of F-actin. All these observations support CD2AP as an essential synaptic regulator, which mutation may contribute to LOAD development.

### How does CD2AP promote spine formation and expansion?

CD2AP is likely a regulator of spine expansion and stabilisation into a functional synapse via F-actin. Notably, CD2AP membrane recruitment is required for actin assembly, which also depends on F-actin(Spence *et al.,* 2016). We demonstrated that CD2AP knockdown decreased spinal F-actin. In contrast, overexpression increased dendritic F-actin, supporting that CD2AP promotes F-actin directly (Lehtonen *et al.,* 2002). CD2AP can cap the barbed ends of F-actin *in vitro,* inhibiting actin polymerisation and depolymerisation (Tang & Brieher, 2013; Wang & Brieher, 2020). The CD2AP capping of actin filaments can also promote branched F-actin dependent on ARP2/3 (Tang & Brieher, 2013), triggering filopodia maturation into spines through spine expansion (Spence *et al.,* 2016). CD2AP-dependent expansion can sequentially recruit PSD-95, functionalising the spine (El-Husseini *et al.,* 2000). In agreement, CD2AP knockdown reduced PSD-95, making fewer synapses and compromising neuronal activity. On the contrary, when CD2AP was in excess, PSD-95 increased in the spines, creating more synapses.

CD2AP could act through its interaction partners, cortactin and capping protein (CP), which have known functions in spines via spinal F-actin regulation (Catarino *et al.,* 2013; Cornelius *et al.,* 2021; Fan *et al.,* 2011; Hering & Sheng, 2003). The CD2AP-cortactin interaction contributes to F-actin stabilisation and accumulation in vivo (Wang & Brieher, 2020). Cortactin knockdown reduces spine density (Catarino *et al.,* 2013; Cornelius *et al.,* 2021; Hering & Sheng, 2003), similar to CD2AP knockdown. Cortactin stabilizes the ARP2/3 dependent branched F-actin, delays its depolymerisation, and can directly activate the ARP2/3 complex (Weaver *et al.,* 2001), supporting a similar role for CD2AP. However, unlike CD2AP, cortactin overexpression leads to longer spines with a minor impact on spine expansion (Hering & Sheng, 2003), suggesting that CD2AP may function in spines independently. CD2AP-CP interaction inhibits CP (Bruck *et al.,* 2006; Takeda *et al.,* 2010; Uruno *et al.,* 2006), which binds actin filaments preventing the addition or loss of actin subunits (Billault-Chaumartin & Martin, 2019), and thus could promote spinal F-actin. However, CP knockdown affects spines differently from CD2AP (Fan *et al.,* 2011), suggesting that CD2AP functions in spines independently. Nevertheless, the complex regulation of F-actin in dendrites (Konietzny *et al.,* 2017) and spines (Honkura *et al.,* 2008) by CD2AP and its interactors will require investigation.

### Impact of a LOAD mutation in CD2AP on spines

The coding mutation in CD2AP (K633R) (rs 116754410) was identified in patients with LOAD (Vardarajan *et al.,* 2015) and recently in children with kidney disease (Nandlal *et al.,* 2022). Although predicted to be pathogenic, it is unknown how K633R affects CD2AP in an AD-relevant way. Our data suggest that the CD2AP (K633R) mutation induces an aberrant gain of dysfunctional CD2AP in spines, increasing their density and volume, likely due to the increased stability of F-actin in spines, with consequent reduction in spine motility that could explain the impaired capacity to establish more synapses (Bonhoeffer & Yuste, 2002). Rescue experiments further support that the CD2AP mutant is dysfunctional since it did not rescue the decreased F-actin and PSD-95 density induced by CD2AP knockdown. It will be necessary to determine the impact of the K633R mutation on endogenous CD2AP and *in vivo* synaptic activity and plasticity to establish how it contributes to AD.

### CD2AP as a risk factor for AD synaptic dysfunction

Carriers of CD2AP coding variants are at higher risk of developing LOAD. Interestingly, while large CD2AP truncations, likely causing CD2AP deletion, cause kidney disease in children, a CD2AP coding mutation was recently associated with kidney disease and mild cognitive decline in adult patients (Tsvetkov *et al.,* 2016). The same mutation has also been found in LOAD (Vardarajan *et al.,* 2015).

Our data suggest that CD2AP coding mutations may only slightly alter CD2AP function, which may enhance AD development with ageing.

Thus, it is crucial to determine their impact on synapses and other disease-relevant mechanisms. The most common variants of LOAD are noncoding and are predicted to alter CD2AP expression; however, it is unknown how CD2AP levels change in the brain. Our data on CD2AP knockdown support the hypothesis that a reduction in CD2AP expression may contribute to synapse loss in AD.

However, if CD2AP levels increase, our data suggest that CD2AP may contribute to the hyperexcitability described in early AD (Vossel *et al.,* 2016). Regarding the LOAD coding variant in CD2AP (K633R), although we observed that it increases spine density and volume, it does not affect synapse density. Instead, it may affect synaptic plasticity, which requires spine expansion to accommodate the increase in synaptic strength that underlies long-term potentiation and memory (Bosch & Hayashi, 2012), a hypothesis supported by our result of reduced spine motility of CD2AP mutant spines.

Concerning the contribution of CD2AP to Aβ-dependent synapse dysfunction in LOAD, we found that the increase in intraneuronal Aβ production induced by CD2AP knockdown in dendrites (Ubelmann *et al.,* 2017) was insufficient to account for the spine loss observed. These results indicate that CD2AP may cause synaptic dysfunction directly and upstream of Aβ production in early AD.

CD2AP may also contribute to the later stages of AD. Indeed, CD2AP has been shown to modulate tau-mediated mechanisms (Shulman *et al.,* 2014) and the integrity of the blood-brain barrier (Cochran *et al.,* 2015) and is associated with cognitive functioning in familial AD (Manzali *et al.,* 2021). Variations in the CD2AP locus are also associated with the burden of neuritic plaques (Shulman *et al.,* 2013). CD2AP-positive neuronal inclusions, resembling neurofibrillary tau tangles, have recently been detected (Camacho *et al.,* 2022).

## Conclusion

CD2AP is a regulator of actin-rich dendritic spines, contributing to spine formation and expansion, mechanisms relevant to synaptic structural plasticity necessary for memory, profoundly affected in AD (Herms & Dorostkar, 2016).

## Acknowledgements

We thank the gift of plasmids to Dr M. Cormont (Univ. Nice), Dr A. Shaw (U. Washington), Dr M. Arpin (Institut Curie), Dr A. Steffen (Helmholtz Centre for Infection Research), and Dr D. Trono (EPFL). We thank M. Pinho and F. Mateus for their technical assistance. We thank L. Almeida for the Excel macros. The authors thank the lab members for their helpful discussions and critical manuscript reading. We thank Dr. S. Marques (NOVA Medical School Animal Facility) and Dr. T. Pereira (NOVA Medical School Microscopy Platform).

This project has received funding from iNOVA4Health—UID/Multi/04462/2019, a program financially supported by Fundação para a Ciência e Tecnologia (FCT)/ Ministério da Educação e Ciência through national funds and co-funded by FEDER under the PT2020 Partnership Agreement); Maratona da Saúde 2016; ALZ AARG-19-618007 Alzheimer’s Association; from the research infrastructure PPBI-POCI-01-0145-FEDER-022122 (FCT and Lisboa2020, under the PORTUGAL2020 agreement -European Regional Development Fund.

CGA has been supported by CEECIND/00410/2017 and CEEC/iNOVA4Health (FCT); FM received an FCT doctoral fellowship (PD/BD/128344/2017). JC is a recipient of an FCT doctoral fellowship (2020.04851.BD).

## Contributions

**Conceptualisation:** Cláudia Guimas Almeida; Farzaneh Mirfakhar, Jorge Castanheira **Methodology:** Farzaneh Mirfakhar, Jorge Castanheira, Cláudia Guimas Almeida, José Ramalho; **Formal Analysis:** Farzaneh Mirfakhar, Cláudia Guimas Almeida, Jorge Castanheira; Raquel Domingues **Investigation:** Cláudia Guimas Almeida; Farzaneh Mirfakhar, Jorge Castanheira, Raquel Domingues and José Ramalho; **Writing - Original Draft:** Farzaneh Mirfakhar. **Writing:** Cláudia Guimas Almeida, Jorge Castanheira; **Visualisation:** Cláudia Guimas Almeida; Jorge Castanheira; **Supervision:** Cláudia Guimas Almeida; **Project administration:** Cláudia Guimas Almeida; **Funding acquisition:** Cláudia Guimas Almeida.

## Methods

### Animals

All animal procedures were performed under the EU recommendations and approved by the NMS-UNL ethical committee (147/2021/CEFCM) and the NOVA Medical School Animal Welfare Body (ORBEA).

### Cell culture, cDNA overexpression, shRNA knockdown, and treatments

Primary neuronal cultures of *Mus musculus* were prepared as previously (Almeida *et al.,* 2005) from the the cortices of wild-type females from wild-type females from embryonic day 16 (E16) and male BALB/c mice. Briefly, E16 brain tissue was dissociated by trypsinization and trituration in Dulbecco’s medium Eagle medium (DMEM, Thermo Fisher Scientific) with 10 % foetal bovine serum (heat-inactivated FBS, Thermo Fisher Scientific). Dissociated neurones plated in DMEM with 10 % FBS on poly-D-lysine (Sigma-Aldrich)-coated 6-well plates (3×10^5^cells/cm^2^), glass coverslips (5×10^4^ cells/cm^2^) After 3-16 h, the media was substituted for Neurobasal supplemented with 2 % B27 (Gibco), or BrainPhys Neuronal Medium (BrainPhys) (Stemcell) supplemented with 2 % Neurocult SM1 (Stemcell) that allows for a more physiological neuronal differentiation and activity (Bardy *et al.,* 2015) and 0.5 % P/S and kept at 37 °C in 5 % CO_2_. Neurones cultured in Neurobasal were used at 21 DIV and BrainPhys at 15 DIV. cDNA transfection and shRNA infection were performed after 12 days *in vitro* (DIV) or 7 DIV for neurones cultured in Neurobasal or BrainPhys media, respectively. For rescue experiments, cDNA transfection was performed with 10 DIV neurones (BrainPhys) after shRNA infection at 7 DIV.

For live cell imaging, primary neurones were grown on 18mm glass bottom plates (FluoroDish, Thermo Fischer Scientific; 3x10^5^ cells/cm^2^). Before imaging, the medium was exchanged for 37 ° C prewarmed imaging medium (120 mM NaCl, 3 mM KCl, 2 mM CaCl2, 2 mM MgCl2, 10 mM glucose, 10 mM HEPES) supplemented with B27.

Neuroblastoma Neuro2a cells (N2a) (ATCCCCL-131) were provided by Z. Lenkei (ESPCI-ParisTech). Cells were cultured in DMEM-Glutamax (Thermo Fisher Scientific) with 10 % FBS (Sigma-Aldrich) at 37 °C and 5 % CO_2_. ShRNA infections were performed 24 h after plating for 72 h.

For cDNA overexpression, primary neurones were lipotransfected with Lipofectamine 2000 (Thermo Fisher Scientific) according to the manufacturer’s instructions with a few modifications, 500 ng of DNA per coverslip and neurones were incubated with lipid-cDNA complexes for 5 min at 37 ° C at 5 % CO_2_, and the medium was replaced with conditioned and fresh BrainPhys media. The following cDNA plasmids were used: p-GFP-C1, Cherry-C1, CD2AP-GFP and CD2AP-myc (Cormont *et al.,* 2003), CD2AP(K633R)-GFP and CD2AP(K633R)-myc were generated by site-directed mutagenesis with the primer 5’GAAATAGCAAAGCTGAAGAAAGCTGTTCTGTTG3’ and 5’CAACAGAACAGCTTTCCTCAGCTTTGCTATTTC3’, mCherry-actin (Koestler *et al.,* 2008).

For shRNA infection, the lentivirus vector with yellow fluorescent tag (YFP) encoding CD2AP shRNA oligonucleotides sh1 (shCD2AP-1; GTGGAACCCTGAACAATAAG) and sh2 (shCD2AP-2, GGAACCAATGAAGATGAACTTACA) (Zhao *et al.,* 2013), or vector encoding non-targeting shRNA (shControl; TR30021, OriGene Technologies) were packed in LV by cotransfection with psPAX2 (gift from Didier Trono; Addgene plasmid # 12260) and pMD2.G (gift from Didier Trono; Addgene plasmid # 12259) psPAX2 and helper plasmids in STAR-Rdpro cells (ECACC 04072117). Lentivirus (LVs) were collected by ultra-speed centrifugation and purified with the Lenti-X concentrator (Takara) according to the manufacturer’s protocol.

For CD2AP knockdown, cells were incubated with LVs expressing shControl, shCD2AP-1, and -2 for 5 minutes at 37 ° C in 5 % CO_2_. The LV media was replaced with conditioned media and fresh BrainPhys.

When indicated, γ-secretase was inhibited with 250 nM γ-secretase inhibitor IX (Calbiochem), while BACE1 was inhibited with 10 µM β-secretase inhibitor compound IV (Merck - 565788) or 0.1 % DMSO (solvent) as a control.

Most experiments were performed with three independent cultures, except when indicated.

### Antibodies and Probes

The following primary antibodies were used: anti-CD2AP (Merck, HPA00326, 1:100 (IF); 1:1000 (WB)); anti-PSD95 (D27E11, Cell Signaling, 3450, 1:200 (IF); 1:2500 (WB)); anti-synapsin (Abcam, ab8, 1:200 (IF); 1:2500 (WB)); anti-tubulin (T5168, Sigma-Aldrich, 1:5000 (WB)); anti-GFP (Sicgen, AB0020, 1:200 (IF)); anti-vGluT1 (Merck, MAB5502, 1:100 (IF)); anti-cortactin (p80/85, Merck, 05-180-I-25UL, 1:200 (IF)); anti-MAP2 (Abcam, ab5392, 1:300 (IF)). The secondary antibodies were conjugated to Alexa-488, -555, and -647 (Invitrogen) or HRP (Bio-Rad). Phalloidin conjugated to Alexa-488, - 555 and -647 (Invitrogen) was used to detect F-actin (1:500).

### Immunofluorescence

Immunofluorescence was performed as previously (Almeida et al., 2005; Ubelmann et al., 2017). Briefly, primary neurones were fixed with 4 % paraformaldehyde/4 % sucrose in PBS 1x for 20 min, permeabilized with 0,3 % Triton-X in PBS 1x for 5 min and blocked with 2 % FBS, 1 % BSA in PBS for 1 h at room temperature (RT) before primary antibody incubation for 16 h at 4 ° C. After washing, secondary antibodies were incubated for 1 h at RT. Coverslips were mounted using Fluoromount-G (Southern Biotech).

### Labelling of active synapses with FM4.64

Primary neurones were incubated with the lipophilic dye FM4.64 (10µM) (Thermo Fisher Scientific) for 90 s in a high potassium saline solution (119 mM NaCl (Enzymatic), 70 mM KCl (Enzymatic), 2 mM CaCl_2_ (Sigma), 2 mM MgCl_2_ (Sigma), 5 mM HEPES (Thermo Fisher Scientific) and 30 mM glucose (NZYtech), washed with a low calcium solution saline solution (150 mM NaCl, 5 mM KCl, 0.2 mM CaCl2, 5 mM MgCl2, 5 mM HEPES and 30 mM Glucose) with 1 µM tetrodotoxin (TTX) (Tocris) for 30 s, fixed for 10 min and mounted.

### Immunoblotting

Cell lysates were prepared using modified RIPA buffer [50 mM Tris-HCl (pH 7.4), 1 % NP-40, 0.25 % sodium deoxycholate, 150 mM NaCl, 1 mM EGTA, and 0.1 % SDS, with protease inhibitor cocktail (PIC, Roche)] as described (Burrinha et al., 2021). Proteins separated by 10 % Tris-glycine SDS–PAGE were transferred to nitrocellulose membranes (GE Healthcare) and processed for immunoblotting using the ECL Prime kit (GE Healthcare). Immunoblot images were captured using a ChemiDoc Gel Imaging System (Bio-Rad) within the linear range and quantified by densitometry using the Analyse gels function in ImageJ.

### Brain synaptosomes preparation

Synaptosomes were prepared from the forebrains (including cortex and hippocampus) of 6-month-old (adult) C57BL/6 mice. All steps were performed at 4 °C. The samples were homogenised with a pestle in ice-cold buffer 1 (0.32 M sucrose (Sigma-Aldrich); 10 mM HEPES; 2 mL PIC (1X); 1mM EDTA, 1× complete PIC tablet; pH 7.4; diluted in ddH_2_O) using 10 ml buffer per gram of tissue. The homogenate was centrifuged (10 min, 1000xg, 4°C) to obtain a pellet containing nuclear fractions (P1) and the postnuclear supernatant (S1). S1 was saved and centrifuged (15 min, 10000 xg, 4 °C) to generate a pellet that contains crude synaptosomes (P2) and a supernatant fraction (S2). P2 was resuspended in 1 ml of buffer 1, saved, and centrifuged (15 min, 10000 xg, 4°C), generating the washed synaptosome fraction (P2’). P2’ was lysed by hypoosmotic shock in ice-cold ddH2O + 10mM HEPES, homogenised with a pestle, and let under head-over-heels rotation at 4°C for 30 min to ensure complete lysis. P2’ was further centrifuged (30 min, 21000 xg, 4 °C) to generate a supernatant that contains crude synaptic vesicles (S3) and a pellet that contains synaptosomal membranes (P3). P3 was resuspended in 100 μl modified RIPA buffer.

### MEA recordings and analysis

Primary neurones were cultured as described above and plated in single-well MEA at a density of 50,000 cells in BrainPhys media per well coated with poly-L-ornithine (Sigma-Aldrich) for 16 h at 37 °C and laminin (Sigma) for 4 h. Primary neurones were transduced with lentivirus expressing shCD2AP-1, shCD2AP-2 or shControl as described. Each MEA well (3Brain Prime HD-MEA) contained 4096 recording electrodes coupled to a ground electrode. MEA recordings of basal neuronal network activity were performed at 15 DIV for 30 s at 37 ° C using the BioCamX recording system and BrainWave v.4.5 software (3Brain).

Spikes were detected with the analysis tool "spike detection" of the BrainWave software. The precise timing spike detection (PTSD) algorithm was used to detect negative spikes, with SD of 8.0, 2.0 ms peak lifetime and 1.0 ms refractory period. For the quantification of active electrodes, we considered the ones with a minimum an average firing rate of 3 spikes/second in percentage of the total (4096). The average firing rate refers to the total number of spikes per 30 s of recording. Spiking frequency refers to the number of spikes detected by each electrode per 30 s recording. The cumulative frequency of the firing rate was calculated using Prism 8.0. Bursts were detected by the "burst detector" analysis tool, with a maximum spike interval of 30 ms and a minimum number of 5 spikes. The burst rate was calculated by dividing the total bursts by 30 s of recording. The number and duration of network bursts, synchronised bursts, spiking frequency of network bursts, interburst intervals, and spiking frequency were calculated using Brainwave v4.

### Fluorescence recovery after photobleaching (FRAP)

To measure F-actin FRAP in spines, primary neurones were transfected with GFP, CD2AP-GFP, CD2AP(K633R)-GFP, and mCherry-actin. GFP. GFP and mCherry-actin were imaged using the LSM980 the 63× NA-1.4 oil Plan-Apochromat objective at 37 ° C, and FRAP was performed using the ZEN (Blue 3.3 edition: Zeiss) bleaching module. ROIs of 12 pixels in diameter were placed on 3-6 spines per dendritic segment. Cells were imaged at 1 frame per 1-4 s in a single plane. After five baseline frames, ROIs were photobleached and imaged for 2 min, allowing recovery to reach a stable plateau. FRAP analysis of mCherry-actin fluorescence intensity in bleached spines was performed as described (Koulouras *et al.,* 2018) using EasyFRAP-web (freely accessible platform-independent; https://easyfrap.vmnet.upatras.gr/). Briefly, fluorescence was background subtracted, corrected for photobleaching and baseline normalised to 100 % for each photobleached spine. FRAP recovery and half-time were calculated by fitting an exponential curve.

### Image acquisition

Images were acquired on an epifluorescence microscope, the Zeiss Imager Z2 system (Zeiss, Oberkochen, Germany) equipped with a 20x air EC Plan-Neofluar objective 63× NA-1.4 oil Plan-Apochromat objective and a Zeiss Axiocam 506 mono camera, and on a confocal microscope with super-resolution, LSM980 – AiryScan 2 (Zeiss) equipped with 63× NA-1.4 oil Plan-Apochromat objective. 3D z-stacks were acquired to enable 3D reconstructions as indicated. The samples were imaged in parallel and using identical acquisition parameters for direct comparison.

### Quantitative Bioimaging Analyses

Image analysis was carried out using Fiji (ImageJ 1.53, https://fiji.sc/), ICY (icy.bioimageanalysis.org (de Chaumont *et al.,* 2012), or IMARIS (version 9.5.0) (https://imaris.oxinst.com/).

For the quantification of spinal enrichment, spinal and dendritic F-actin, ICY was used to outline a region of interest (ROI) corresponding to a dendritic section manually or automatically based on the GFP signal and outline spines, either by selecting spines with the "ellipse" selection tool or subtracting the dendritic shaft from the dendritic ROI. Each ROI’s fluorescence mean intensity and area were obtained with ICY ROI measurements. Spinal enrichment was obtained by dividing the fluorescence intensity in the spine ROI by the fluorescence intensity in the dendritic ROI and presented as a percentage. Spinal F-actin puncta was segmented using the ICY "Spot detector" module in the spines ROI, and each puncta area was obtained using ICY ROI measurements.

For the quantification of vGluT1, PSD-95 and FM4.64 puncta density or size, dendritic ROIs were outlined with the ICY ’Area’ selection tool, puncta were segmented with the ICY "Spot detector" module and ROI’s measurements were exported as previously including the dendrite ROI length (Feret’s diameter) (Burrinha *et al.,* 2021). Alternatively, dendritic ROIs were outlined using the ’polygon selection’ tool on FIJI, and the "ComDet v.0.5.5" plugin was used for segmentation, colocalisation considering the max distance between objects less than 5 pixels, and the other measurements as previously (Burrinha *et al.,* 2023).

For the Sholl analysis, single pyramidal neurones expressing GFP were 3D reconstructed using IMARIS’ Filament Tracer. The Filament Sholl analysis was used to quantify neurite number, intersections, and length.

To quantify spine density volume, IMARIS "Filament Tracer" was used to 3D reconstruct dendrites based on the GFP (shRNA) or mCherry (OE) signal. The spine module automatically detects spines and provides spine head volume. When automatic detection was impossible, spine density was manually counted using the point selection tool in FIJI.

To quantify spine motility, dendrites were 4D reconstructed, and the spine volume was measured using IMARIS in dendrites imaged live 1 frame every 30 s for 30 min. The spine motility index was measured as described (Dunaevsky *et al.,* 1999) with slight modifications. Briefly, the motility index was obtained for each spine by calculating the ratio of the difference between the largest and smallest spine volume to the average spine volume during the time-lapse.

### Statistics

Prism 8 (GraphPad) was used for statistical analysis and for graphic representation of individual or average replicates with mean ± S.D. as indicated in the figure legends. The sample size was determined on the basis of pilot studies. Data were tested with the D’Agostino-Pearson omnibus normality test. For paired data, the Wilcoxon t-test was applied. For nonparametric and unpaired data, the Mann–Whitney test was applied. For parametric and multiple comparisons, statistical analysis of data ordinary one-way ANOVA with Holm-Sidlak’s multiple comparisons test was applied. For nonparametric and multiple comparison statistical data analysis, one-way ANOVA (Kruskal-Wallis test) with post hoc Dunn’s testing was applied. Significance was considered as \**P*<0.05, \*\**P*<0.01, \*\*\**P*<0.001, \*\*\*\**P*<0.0001; ns, not significant.

---

**Figure S1.**
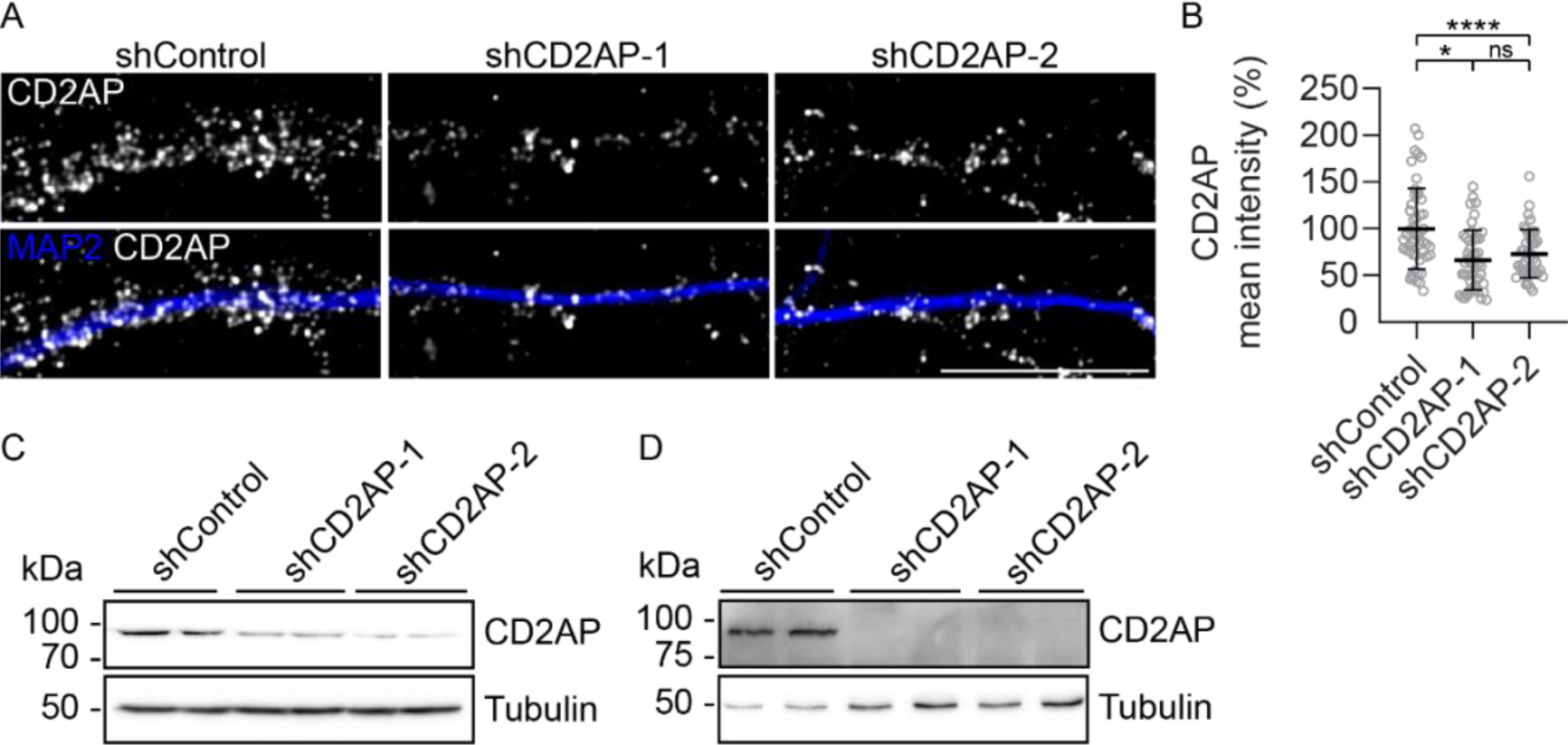
Knockdown of CD2AP by treatment with lentivirus expressing CD2AP shRNA. **(A)** CD2AP (grey) and MAP2 (blue) in dendrites of neurones treated with shControl, shCD2AP-1 and shCD2AP-2. Scale bar: 10 µm. **(B)** Quantification of CD2AP mean intensity in dendrites (n=3, N_shControl_=50, N_shCD2AP-1_=46, N_shCD2AP-2_=42). **(C-D)** CD2AP western blot and tubulin as loading control in Neuro2a cells **(C)** and puromycin-selected NIH 3T3 cells **(D)** treated with shControl, shCD2AP-1 and shCD2AP-2.

**Figure S2.**
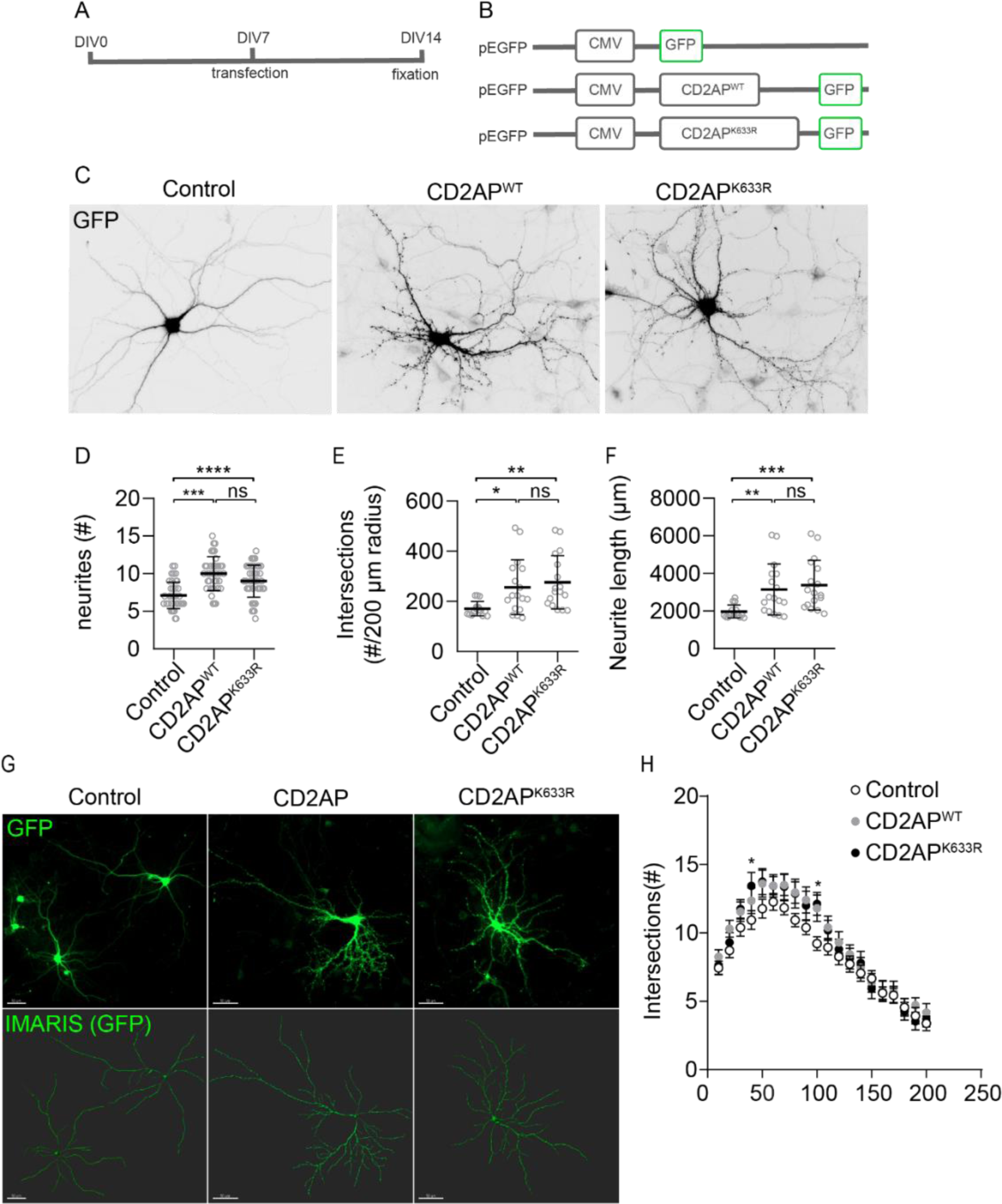
CD2AP^WT^, but not CD2AP^K633R^, overexpression increases neurite number, length, and branching. **(A)** Overexpression experiment timeline indicating the day *in vitro* when neurones are transfected and fixed. **(B)** cDNA plasmids used encoding GFP, CD2AP^WT^-GFP, and CD2AP^K633R^-GFP. **(C)** Neurones overexpressing GFP, CD2AP^WT^, and CD2AP^K633R^ (grey). **(D)** Quantification of neurites number in neurones expressing GFP, CD2AP^WT^, and CD2AP^K633R^ (n=6, N_GFP_=42, N_CD2AP_^WT^=37, N ^K633R^=46). **(E)** Quantification of the number of intersections per 200 µm radius in neurones expressing GFP, CD2AP^WT^, and CD2AP^K633R^ (n=3, N_GFP_=15, N_CD2AP_^WT^=17, N ^K633R^=17). **(F)** Quantification of neurite length in neurones expressing GFP, CD2AP^WT^, and CD2AP^K633R^ (n=3, N_GFP_=15, N_CD2AP_^WT^=17, N ^K633R^=17). **(G)** Low magnification (20x) images of neurones expressing GFP, CD2AP^WT^, and CD2AP^K633R^ (green) and respective 3D IMARIS reconstructions. Scale bar: 50 µm. **(H)** Sholl analysis of the number of intersections of neurones expressing GFP, CD2AP^WT^, and CD2AP^K633R^ (n=3, N_GFP_=18, N_CD2AP_^WT^=20, N_CD2AP_^K633R^=16). Data are presented as mean ± SD. \**P*<0.05, \*\**P*<0.01, \*\*\**P*<0.001, \*\*\*\**P*<0.0001; ns, not significant.

**Figure S3.**
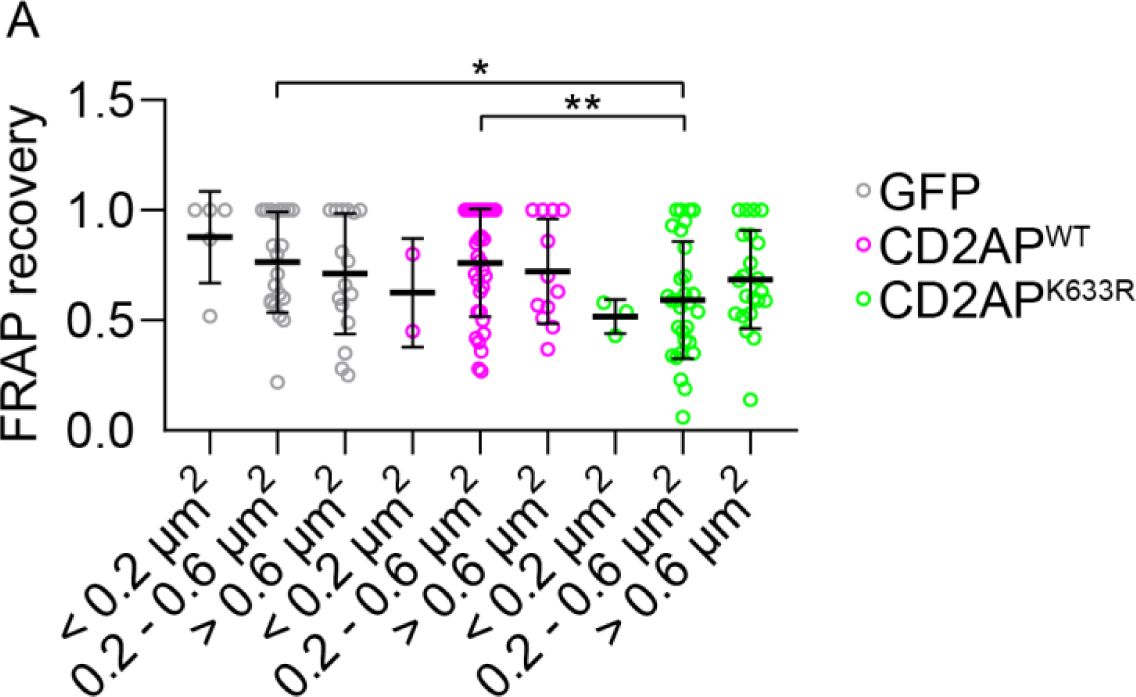
CD2AP^K633R^ mutation decreases the mCherry-actin FRAP recovery, especially in the intermediate spine size. **(A)** Quantification of FRAP time recovery of mCherry-actin in spines expressing GFP, CD2AP^WT^ and CD2AP^K633R^ per size category: small (<0.2 µm^2^), intermediate (0.2-0.6 µm^2^) and large (>0.6 µm^2^) (n=3, N_GFP_(<0.2)_=_5, N_GFP_(0.2-0.6)=21, N_GFP_(>0.6)=16, N_CD2AP_^WT^(<0.2)=2, N_CD2AP_^WT^(0.2-0.6)=34, N_CD2AP_^WT^(>0.6)=12, N_CD2AP_^K633R^(<0.2)=3, N_CD2AP_^K633R^(0.2-0.6)=30, N_CD2AP_^K633R^(>0.6)=22 spines). Data are presented as mean ± SEM. \**P*<0.05, \*\**P*<0.01.

